# Laterality of the frontal aslant tract (FAT) explains externalizing behaviors through its association with executive function

**DOI:** 10.1101/162495

**Authors:** Dea Garic, Iris Broce, Paulo Graziano, Aaron Mattfeld, Anthony Steven Dick

**Affiliations:** Department of Psychology, Florida International University, Miami, FL, 33199; Department of Radiology and Biomedical Imaging, University of California, San Francisco, San Francisco, CA, 94143

## Abstract

We investigated the development of a recently-identified white matter pathway, the frontal aslant tract (FAT) and its association to executive function and externalizing behaviors in a sample of 129 neurotypical children ranging in age from 7 months to 19-years. We found that the FAT could be tracked in 92% of those children, and that the pathway showed age-related differences into adulthood. The change in white matter microstructure was very rapid until about 6-years, and then plateaued, only to show age-related increases again after the age of 11-years. In a subset of those children (5-18-years; *n* = 70), left laterality of the microstructural properties of the FAT was associated with greater attention problems as measured by the Child Behavior Checklist (CBCL). However, this relationship was fully mediated by higher executive dysfunction as measured by the Behavior Rating Inventory of Executive Function (BRIEF). This relationship was specific to the FAT—we found no relationship between laterality of the white matter of the brain in general and attention problems, or executive function. These findings suggest that the degree to which the developing brain favors a right lateralized structural dominance of the FAT is directly associated with executive function and attention. This novel finding provides a new potential structural biomarker to assess attention deficit hyperactivity disorder (ADHD) and associated executive dysfunction during development.

**Significance Statement:** To investigate the function of a recently-identified white matter fiber pathway, the frontal aslant tract (FAT), we tracked the pathway in 129 typically developing children using diffusion-weighted magnetic resonance imaging (DW-MRI). We then examined whether laterality of the tract is associated with attention problems and executive function. We found that reduced right laterality of the tract was associated with greater executive dysfunction, which predicted increased reports of attention problems. The findings suggest that the degree to which the developing brain favors a right lateralized structural dominance of the FAT is directly associated with executive function and attention. This novel finding provides a new potential structural biomarker to assess attention deficit hyperactivity disorder (ADHD) and associated executive dysfunction during development.

---

It is critical to understand the etiology of children’s early externalizing behavior problems, including symptoms of Attention-Deficit/Hyperactivity Disorder (ADHD) such as inattention, hyperactivity, and impulsivity. These are the most common reason for early childhood mental health referrals (Keenan and Wakschlag, 2000; Thomas and Guskin, 2001) and occur in 10-25% of preschoolers (Carter et al., 2004; Furniss et al., 2006). Despite successful development of evidence-based treatments for ADHD, early interventions have little impact on children’s long-term academic and social impairment (Jensen et al., 2007; Molina et al., 2009; Molina et al., 2013; Sonuga-Barke et al., 2013). Researchers, clinicians, and patients are thus desperate for tangible progress in identifying biomarkers for treatment of mental illness in both adults and children. Identifiable biomarkers can serve as indicators of treatment response, as indicators of heterogeneity within broadly defined disorders, or as future targets of non-invasive brain stimulation treatments, and are necessary for applying precision medicine approaches to mental health treatment.

Here we investigate a recently-identified white matter fiber pathway, the frontal aslant tract (FAT), and attempt to define its functional relevance to executive function and externalizing behaviors— namely, attention problems—in a sample of typically developing children. The function of the FAT remains a matter of speculation, and its investigation in children has been minimal (Madsen et al., 2010; Broce et al., 2015). Based on the fiber pathway’s putative connectivity joining the posterior inferior frontal gyrus (IFG) with the pre-supplementary and supplementary motor areas (pre-SMA and SMA; Catani et al., 2012; Kinoshita et al., 2012; Martino and De Lucas, 2014; Bozkurt et al., 2016; Szmuda et al., 2017), investigators have focused on the pathway’s involvement in speech and language function. For example, stimulation of the left FAT during awake surgery induces speech arrest (Vassal et al., 2014; Fujii et al., 2015; Kinoshita et al., 2015), and the left FAT is associated with executive control of speech and language in other tasks (e.g., verbal fluency, stuttering; Catani et al., 2013; Basilakos et al., 2014; Mandelli et al., 2014; Broce et al., 2015; Kinoshita et al., 2015; Sierpowska et al., 2015; Kemerdere et al., 2016; Kronfeld-Duenias et al., 2016).

However, given the well-known laterality of function in the brain (Toga and Thompson, 2003; Herve et al., 2013), the possibility remains that the function of the left FAT differs from its homologue on the right. Indeed, Aron et al (2014) suggested that the right posterior IFG, the pre-SMA, and the connections between those regions (i.e., via the FAT) are associated with inhibitory control in executive function tasks (Aron et al., 2007), a possibility supported by fMRI, electrocorticography (ECoG), and diffusion-weighted imaging (DWI) data in adults (Swann et al., 2012). It is thus possible that while the left FAT might be associated with executive control of speech and language function (e.g., in the case of verbal fluency or speech initiation), the right FAT might be associated with executive control of action (e.g., inhibitory control of action). Consistent with this proposition, functional imaging data suggest that lateralization of these functions emerges during childhood (Holland et al., 2001; Everts et al., 2009). Furthermore, ADHD is associated with structural and functional abnormalities in the pre-SMA and right IFG regions connected by the FAT (Rubia et al., 1999; Mostofsky et al., 2002; Suskauer et al., 2008a; Suskauer et al., 2008b). However, the direct contribution of the FAT to executive function, or to externalizing behaviors more broadly, during development has not been investigated.

We explored this issue in a DWI study of neurotypical children between the ages of 7-months and 19-years. We tracked the left and right FAT in these participants and related diffusion metrics of white matter microstructure to behavioral inventories of executive function, and attention. Based on the right-lateralized associations with IFG and pre-SMA function and executive function, we predicted that deviation from right lateralization of this pathway would be associated with poorer executive function, and increased instances of externalizing behaviors.

## Materials and Methods

### Participants

In the present study, we analyzed a publically available data set of neurotypical children from the Cincinnati MR Imaging of NeuroDevelopment (C-MIND) database, provided by the Pediatric Functional Neuroimaging Research Network (https://research.cchmc.org/c-mind/) and supported by a contract from the Eunice Kennedy Shriver National Institute of Child Health and Human Development (HHSN275200900018C). The data are available from CMIND by request, which facilitates validation of the results we report here. Participants in the database are full-term gestation, healthy, right-handed, native English speakers, without contraindication to MRI. By design, the C-MIND cohort is demographically diverse (38% nonwhite, 55% female, median household income $42,500), intended to reflect the US population.

We tracked the FAT in all available participants (*n* = 129; 70 females). The age range for the full sample was 7-months to 19-years (*M* = 8.8 years; *SD* = 5.0 years). From the full sample, 70 participants had behavioral data on all of the measures of interest, and also had the tracked fiber pathways of interest. Thus, the sample size for the mediation analysis we report below is *n* = 70. In this subset, the participants were equally split by gender (35 females), and ranged in age from 5-years to 18-years (*M* = 10.9 years; *SD* = 3.7 years). A wide range was represented on the measure of socioeconomic status, which was coded on a 10-point ordinal scale of household income (‘0’ = $0 - $5000 to ‘10’ = Greater than $150,000; *M* = 5.1; *SD* = 2.6). In the subsample, all of the children were typically developing and the sample was made up of 94% Non-Hispanic/Non-Latino participants. The study was approved by the Cincinnati Children’s Hospital Medical Center Institutional Review Board. The Florida International University Institutional Review Board approved the data use agreement.

### Experimental Design and Statistical Analysis

We employed analysis of a quasi-experimental design on a publically available dataset consisting of DWI MRI scans, and parent/teacher report measures of executive function (i.e., the Behavior Rating Inventory of Executive Function; BRIEF) and externalizing behaviors (focusing on Attention Problems with the Child Behavior Checklist; CBCL). We conducted High Angular Resolution Diffusion Imaging (HARDI)-based analysis of the DWI data using a generalized q-sampling imaging (GQI) model-free reconstruction method (Yeh et al., 2010). We manually reconstructed the FAT in each hemisphere of each subject, defined on the original image space of the subject. We then explored the age-related change in the pathway’s microstructure, and calculated laterality of the pathway. Following that, we conducted a simple mediation analysis in which laterality of the FAT was entered as a predictor, executive function as measured by the BRIEF was entered as a mediator, and CBCL Attention Problems was entered as the outcome. The details of these steps are presented below.

### MRI Scans

Single shell, 61 direction HARDI scans were created using a spin-echo, EPI method with intravoxel incoherent motion imaging (IVIM) gradients for diffusion weighting of the scans. They were acquired using a 32-channel head coil (SENSE factor of 3), which obtained 2 × 2 × 2 mm spatial resolution at *b* = 3000 (EPI factor = 38, 1752.6 Hz EPI bandwidth, 2 × 2.05 × 2 acquisition voxel; 2 × 2 × 2 reconstructed voxel; 112 × 109 acquisition matrix). The scan took under 12 minutes, with an average scan time of 11 minutes and 34 seconds. Seven *b* = 0 images were also acquired at intervals of 8 images apart in the diffusion direction vector. These *b*0 images are used for co-registration and averaged to form the baseline for computation of the diffusion metrics of interest.

#### HARDI post-processing

The image quality of the HARDI data was assessed using DTIPrep (http://www.nitrc.org/projects/dtiprep), which discards directions as a result of slice dropout artifacts, slice interlace artifacts, and/or excessive motion. Participants with 45 directions or more were included in the study (Tournier et al., 2013). All usable data were registered to the reference image (*b* = 0), using a rigid body mutual information algorithm and were eddy current corrected for distortion. Because the age of the participants ranges from 7 months to 19-years, we did not warp the images to a standard template, and instead defined the pathways of interest in the original diffusion image space.

Using DSI Studio, we used the GQI model-free reconstruction method, which quantifies the density of diffusing water at different orientations (Yeh et al., 2010) to reconstruct the diffusion orientation distribution function (ODF), with a regularization parameter equal to 0.006 (Descoteaux et al., 2007). From this, we obtained normalized Quantitative Anisotropy (nQA).

QA is defined as the amount of anisotropic spins that diffuse along a fiber orientation, and it is given mathematically by:

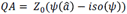

where *ψ* is the spin distribution function (SDF) estimated using the generalized q-sampling imaging, â is the orientation of the fiber of interest, and *iso(*() is the isotropic background diffusion of the SDF. Z_0_ is a scaling constant that scales free water diffusion to 1 (i.e., it is scaled to the maximum ODF of all voxels, typically found in cerebral spinal fluid). This allows the QA value will mean the same thing across different participants (Yeh et al., 2010).

QA can be defined for each peak in the SDF. Because deterministic tractography (which we use in this study) follows individual peaks across a streamline of voxels, we have focused on the first peak (QA_0_). Unlike typical diffusion-tensor imaging (DTI) metrics such as FA, QA must be further normalized so that it can be compared across different participants. This normalized QA metric, nQA, was calculated according to the generalized *q*-sampling imaging method described above (Yeh et al., 2010), and essentially normalizes the maximum QA value to 1. GQI performs as well as other HARDI metrics, such as Constrained Super-resolved Spherical Deconvolution (CSD; Tournier et al., 2007; Yeh et al., 2013)) and better than standard DTI algorithms (Yeh et al., 2013; Daducci et al., 2014).

In summary, we used the GQI reconstruction to map the streamlines, with deterministic tractography following the QA_0_ at each voxel. We used the nQA_0_ component in our analysis of the relation of white matter microstructure to behavior. To facilitate comparisons with prior literature, we report the DTI FA metric for assessment of age-related differences.

#### Defining the FAT: Drawing the Regions of Interest (ROIs)

There were eight total ROIs manually drawn for each participant, four per hemisphere. In each hemisphere, we drew ROIs for two superior frontal gyri: the pre-SMA and SMA; and two inferior frontal gyri ROIs, the IFGOp and the IFGTr.

We systematically drew the eight ROIs for each participant following the same sequence of steps, starting from identification of the brain’s midline slice. From the midline slice, the anterior commissure was located, which represents the arbitrary dividing line between the pre-SMA and SMA ROIs (Kim et al., 2010; Vergani et al., 2014). The superior border for both ROIs is the top of the brain, and the inferior border is the cingulate gyrus. The pre-SMA ROI’s anterior border is the anterior tip of the cingulate while the posterior border for the SMA is the precentral sulcus. The inferior frontal gyri ROIs—*pars triangularis* (IFGTr) and *pars opercularis* (IFGOp)—were parcellated with reference to the Duvernoy atlas (Duvernoy et al., 1999), and were defined by the semi-automated Freesurfer parcellation (Desikan et al., 2006). All ROIs were visually inspected and edited to include the underlying white matter. Fiber tracking was terminated when the relative QA for the incoming direction dropped below a preset threshold (0.02-0.06, depending on the subject; Yeh et al., 2010) or exceeded a turning angle of 40°.

#### Calculation of Laterality

We calculated FAT laterality (*L*) following the standard formula (Thiebaut de Schotten et al., 2011):

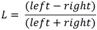

The HARDI metric nQA_0_ was used as the measure of interest. Thus, positive values indicate greater left laterality.

### Behavioral Measures

#### Behavior Rating Inventory of Executive Function (BRIEF)

The Behavior Rating Inventory of Executive Function (BRIEF; Gioia et al., 2000) was used to assess executive function. The BRIEF is a parent- and teacher- report measure of executive function. It has eight subscales, which have been grouped, based on factor analysis of these scales, into two indices, the Metacognitive Index (MI) and the Behavioral Regulation Index (BRI). The BRI is comprised of the Inhibit, Shift, and Emotional Control subscales, and reflects the ability to set shift and control behavior through the administration of appropriate inhibitory control. The MI is comprised by the Initiate, Working Memory, Plan/Organize, Organization of Materials, and Monitor subscales. This index assesses the ability to initiate, plan, and organize behavior, and to apply and sustain appropriate working memory to control behavior (Gioia et al., 2002). All eight subscales comprise a Global Executive Composite (GEC) score. The BRIEF has clinical utility for the diagnosis of ADHD (Isquith and Gioia, 2000). For example, McCandless and O’Laughlin (2007) found that the MI was sensitive to the diagnosis of ADHD, while the BRI was most sensitive to dissociating among subtypes of ADHD. The MI, BRI, and GEC composite scores were the focus of the present investigation.

#### Child Behavior Checklist (CBCL)

The Child Behavior Checklist (CBCL; Achenbach and Rescorla, 2000, 2001, 2003) was administered using either the preschool, school-age, or adult form (depending on the participant’s age). We focused on the Attention Problems outcome scale, which has high reliability (*r* = 0.78 for the preschool form; *r* = 0.92 for the school age form, with *r* = 0.70 and 0.60 for 12- and 24-month follow-up, respectively; *r* = 0.87 for the adult form). This scale is also highly associated with ADHD diagnosis (Biederman et al., 1993; Papachristou et al., 2016).

### Simple Mediation Analysis

We examined the relationships among the laterality of the FAT, executive function, and attention in typical individuals using a simple mediation model. This model was statistically analyzed in SPSS v23 within the PROCESS regression framework from Hayes (Hayes, 2013). We used Model 4 in the framework. Three mediation models were tested. In the first model, we tested whether the BRIEF GEC—which includes all subtests of the BRIEF—mediates the relation between laterality of the FAT and Attention Problems. Because some of the ratings on the BRIEF are directly related to items on the CBCL Attention Problems subscale (e.g., “Impulsive or acts without thinking”), we re-ran the same analysis replacing GEC with MI as a mediator, which mitigates that potential confound. Although not completely orthogonal, we also ran the analysis with BRI as the mediator. In the mediation analysis, the following covariates were included: gender, age (in days), whole brain nQA (to control for general white matter microstructure), and household income (on a 10-point scale, to control for SES). Because these controls were included, raw scores were used for the outcome variables. In addition, to confirm whether the results we report were specific to the FAT, we also ran the same mediation model with laterality of the whole brain white matter as the predictor of interest.

## Results

### Identification of the Fiber Tracts

Using the individually defined ROIs, we were able to track four subcomponents of the FAT, in the following percentage of participants from the full sample (*n* = 129; averaged across the hemispheres; Figure 1): IFGOp <-> pre-SMA (92%); IFGTr <-> pre-SMA (66%); IFGOp <-> SMA (76%); IFGTr <-> SMA (26%). However, the largest component defined the connections between the IFGOp and pre-SMA, and this was tracked in almost all participants for both hemispheres. This replicates the pattern of connectivity reported in adults (Catani et al., 2012; Kinoshita et al., 2012; Martino and De Lucas, 2014; Bozkurt et al., 2016; Szmuda et al., 2017). Furthermore, components overlap to a significant degree as they traverse the frontal white matter, and thus analysis of these components introduces a dependency in the results. Finally, the available literature suggests that the IFGOp and pre-SMA are most likely to be associated with executive function (Aron et al., 2007; Swann et al., 2012). Therefore, for analytic and conceptual simplicity, we focused on the IFGOp <-> pre-SMA component for the age-related and mediation analyses described below.

**Figure 1.**
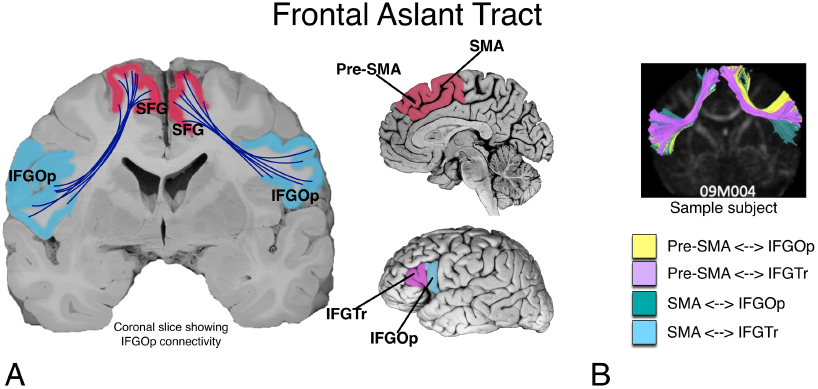
Illustration of the putative connectivity of the frontal aslant tract (FAT). A. Connectivity of the tract is bilateral between the inferior frontal gyrus (*pars opercularis* (IFGOp) and *pars triangularis* (IFGTr) and the superior frontal gyrus (namely, pre-supplementary motor area (pre-SMA) and supplementary motor area (SMA)). B. The pathway can be further differentiated into four parts connecting two parts of the IFG to the pre-SMA and SMA.

### Age-related Differences in Fractional Anisotropy

In Figure 2 we show the age-related differences in FA of the left (purple) and right FAT (in teal; IFGOp <-> pre-SMA component). These are mapped along with the general trend of white matter development in the whole brain (grey line). Shading represents the 95% confidence intervals.

**Figure 2.**
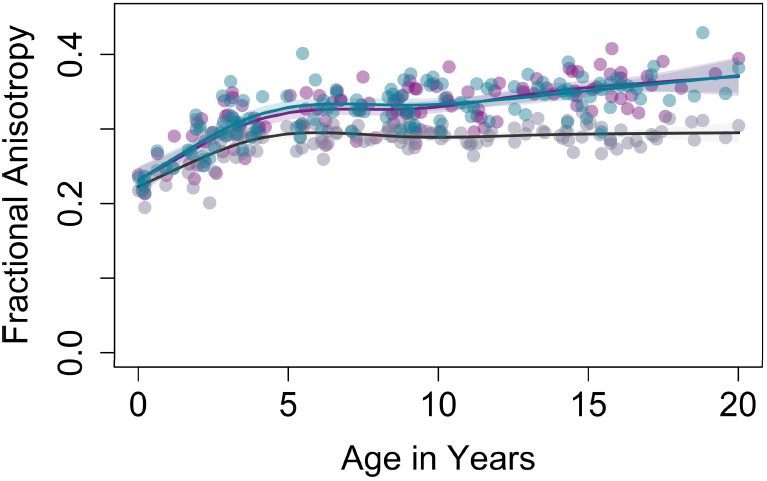
Age-related differences in FA of the left (purple) and right FAT (in teal; IFGOp <-> pre-SMA component). These are mapped along with the general trend of white matter development in the whole brain (grey). Shading represents the 95% confidence intervals.

Visual inspection of the scatter plots indicated that the data might be summarized by a non-linear model. Using the R splines package (v. 3.2.3) to fit restricted cubic splines (package *rms*), and fitting robust models (package *rlm*), we fit two models for each dependent measure. The first was a linear model of age and gender predicting FA. The second used restricted cubic spline regression (specifying two knots), plus gender, as predictors. However, there were no effects of gender in any of the models (smallest *p* = 0.26), so that variable was dropped. Compared to the linear models, AIC and BIC fit indices were larger for the non-linear models (Left FAT Linear: AIC = −512.8; BIC = −504.5; Non-linear: AIC = −534.4; BIC = −517.7; Right FAT Linear: AIC = −509.1; BIC = −500.8; Non-linear: AIC = −540.9; BIC = −524.3; Whole brain Linear: AIC = −635.2; BIC = −626.6; Non-linear: AIC = −697.4; BIC = −680.2) Thus, the non-linear models are reported here, and were significant for the left FAT (*F*(4, 114) = 47.1, *p* < .001), the right FAT (*F*(4, 114) = 44.4, *p* < .001), and the whole brain (*F*(4, 124) = 36.89, *p* < .001). Inspection of the plot reveals that age-related differences in white matter appear rapidly over the first 6-years. However, while for the whole brain the differences in white matter plateau, for the FAT there is subsequent increase in FA after about age 11.

### Mediation Analysis

The age-related differences in white matter suggest that age might be a potential “third variable” driving the association between white matter and behavior. Therefore, we first explored whether age was associated with the behavioral scores. It was not for BRIEF GEC or MI (*r* = −0.16 for BRIEF GEC, *t*(68) = 1.31, *p* = 0.20; *r* = −0.14 for BRIEF MI, *t*(68) = 1.18, *p* = 0.24) or for CBCL Attention Problems (*r* = −0.10, *t*(68) = 0.80, *p* = 0.42). However, there was a small correlation between age and BRIEF BRI scores (*r* = −0.27, *t*(68) = 2.31, *p* = 0.03). To mitigate this possible confound, we controlled for age and other covariates (sex, whole brain white matter, and SES) in the analysis. With the exception of the relation between age and BRIEF BRI, no significant effects for the covariates were found, and the findings are reported with the covariates included in the model. This suggests that the mediation analysis speaks to individual differences in the measures of FAT white matter correlates that predict differences in executive function and externalizing behaviors.

Figure 3 and Table 1 show the results of the mediation analysis. Specifically, the results show that left laterality of the FAT predicted higher CBCL Attention Problems scores. It also predicted higher BRIEF scores. Thus, when considered in a mediation model, the relation between FAT laterality and Attention Problems was fully mediated by executive function as measured by the BRIEF. This finding held for the full executive function index of the BRIEF (i.e., GEC), and also for the MI and BRI considered separately. Because higher scores on each of the outcome variables reflect greater executive function and attention problems, our analysis shows that greater left laterality predicts more executive dysfunction, and higher reports of attention problems.

**Figure 3.**
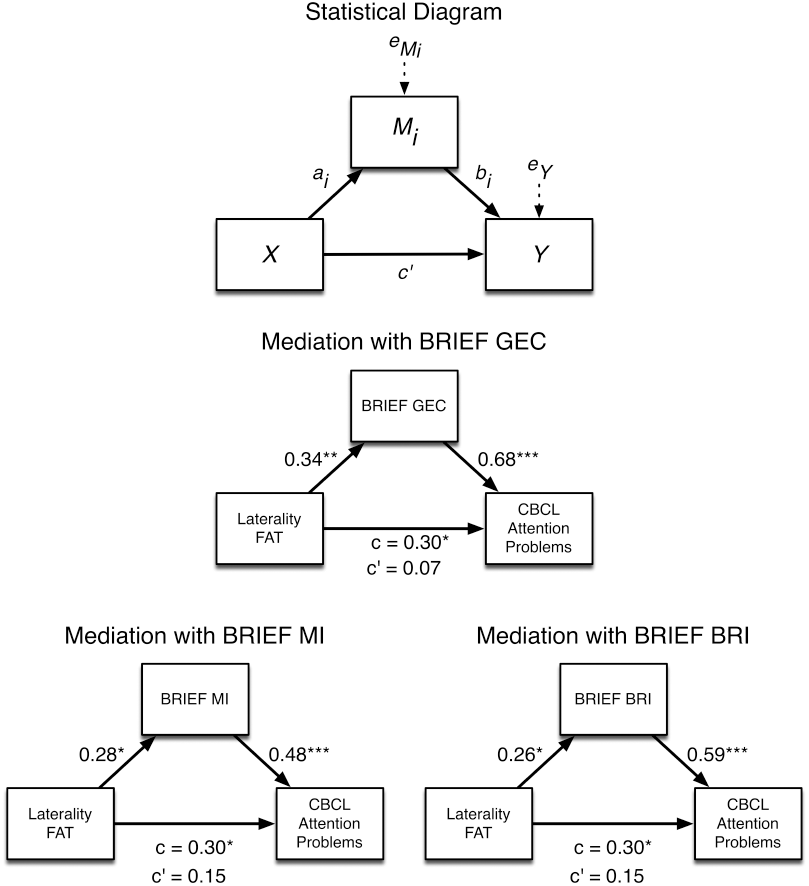
Mediation analysis showing greater left laterality of the FAT predicts more executive dysfunction on the BRIEF, and higher reports of attention problems on the CBCL. The statistical model is presented (top), and is tested with BRIEF GEC as the mediator (middle), BRIEF MI as the mediator (bottom left), and BRIEF BRI as the mediator (bottom right).

**Table 1.**
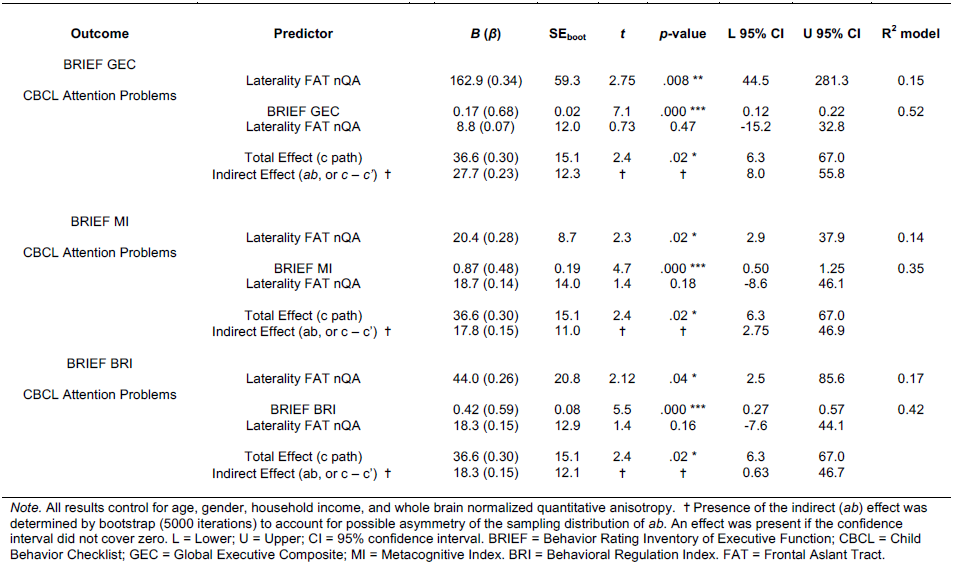
Results of the Mediation Analyses for Frontal Aslant Tract laterality predicting Attention Problems

This pattern of results was not apparent when we assessed laterality of the whole brain white matter. Laterality of the whole brain white matter did not predict CBCL Attention Problems (*B* = 19.7, *t*(64) = 0.36, *p* = 0.72, 95% CI = −88.4 to 127.7), nor did it predict BRIEF GEC (*B* = 96.3, *t*(64) = 0.45, *p* = 0.66, 95% CI = −332.3 to 525.0), BRIEF MI (*B* = 28.8, *t*(64) = 0.93, *p* = 0.36, 95% CI = −33.2 to 90.8), or BRIEF BRI (*B* = 70.4, *t*(64) = 0.96, *p* = 0.34, 95% CI = −76.0 to 216.7). There was no mediation effect (the 95% CI for the *ab* parameter covered zero for any models; *B* = 16.9 (−65.3 to 87.0) for BRIEF GEC, *B* = 27.3 (−32.5 to 91.5) for BRIEF MI, and *B* = 31.3 (−32.8 to 102.5) for BRIEF BRI). This suggests that the finding we report is specific to the FAT.

## Discussion

We investigated the development of the FAT and its association to executive function and externalizing behaviors in a sample of 129 children ranging in age from 7-months to 19-years. We found that the FAT could be tracked in over 90% of those children, and that the pathway showed age-related differences into adulthood. The change in white matter microstructure was very rapid until about 6-years, and then plateaued, only to show age-related increases again after the age of 11-years. In a subset of those children for whom behavioral data was available (5-18-years; *n* = 70), left laterality of the microstructural properties of the FAT predicted greater attention problems as measured by the CBCL. However, this relationship was fully mediated by higher executive dysfunction as measured by the BRIEF. This relationship was specific to the FAT—we found no relationship between laterality of the white matter of the brain in general and attention problems, or executive function. These findings suggest that the degree to which the developing brain favors a right lateralized structural dominance of the FAT is directly associated with developing executive function and attention. This novel finding provides a new potential structural biomarker for attention problems and associated executive dysfunction, which would lay the foundation for future exploration as a biological indicator of treatment response in developmental externalizing disorders, such as ADHD.

### The Role of the Frontal Aslant Tract in Executive Function

Our findings are consistent with current neurobiological models of executive function in adults. For example, several authors (Aron et al., 2007; Wiecki and Frank, 2013; Aron et al., 2014; Jahanshahi et al., 2015; Aron et al., 2016) have proposed a model for stopping behavior—i.e., countermanding an initiated response tendency via top-down executive control, recruited during Go/NoGo and Stop-Signal experimental paradigms. In these tasks, a prepotent response is initiated (a Go process) that must be over-ridden when a stop-signal occurs (the Stop process). These models propose that stopping requires the integrity of the right IFG and the pre-SMA, and that these regions form part of a cortico-basal ganglia “network for inhibition” (Jahanshahi et al., 2015).

In our study, we replicate the structural connection of the IFG and pre-SMA via the fibers of the FAT, and other research confirms the functional connectivity of these two regions (Duann et al., 2009). The establishment of this monosynaptic connection between the IFG and pre-SMA is important for exploring the distinct roles each region plays within this “network for inhibition” (Ridderinkhof et al., 2004). In the Wiecki/Frank computational model (Wiecki and Frank, 2013), the right IFG directly activates neurons of the subthalamic nucleus, which plays an explicit role in stopping motor behavior (Favre et al., 2013; Jahanshahi, 2013; Obeso et al., 2014; van Wouwe et al., 2017). However, others suggest that this connection may proceed via the pre-SMA (Aron et al., 2016). This is important to work out, and our results suggest that the connection between IFG and pre-SMA is an important structural component of this network. Further, it may be that the modulation of subthalamic nucleus activity proceeds through this link. From this perspective, the right FAT is a pathway for inhibition. Indeed, higher FA in the white matter under the pre-SMA and right IFG is associated with better response inhibition in children (Madsen et al., 2010) and older adults (Coxon et al., 2012).

It is also possible, though, that the pathway does not play a role in inhibition, but rather in conflict detection. Thus, the pre-SMA may be critical for the detection of and resolution of conflicting action representations—the “winning” representation is reinforced, and the “losing” representation is suppressed (Nachev et al., 2007). Indeed, Nachev and colleagues (2007) found that a patient with a focal lesion to the right pre-SMA was significantly impaired on a task requiring the resolution of conflict between competing action plans. This is consistent with fMRI task paradigms showing activation of right pre-SMA in situations in which a participant must choose to perform a new response in favor of an established response (Garavan et al., 2003), and in single-unit recording of a human in which pre-SMA neurons appear to play a role in the selection and preparation of movements (Amador and Fried, 2004). The pre-SMA and its connections with the IFG appear to be important for these processes.

### The Role of the Frontal Aslant Tract in Externalizing Behaviors and Attention

We also showed that microstructural properties of the FAT, as measured by DWI, are associated with increased reports of attention problems in children. This is consistent with prior neuroimaging research in people with ADHD showing activation differences compared to neurotypical people in right IFG and pre-SMA during executive function tasks. For example, people with ADHD show hypoactivation of the right IFG during Go/NoGo and SST tasks (Rubia et al., 1999). Anatomic and functional differences in children with ADHD are also reported for the pre-SMA (Mostofsky et al., 2002; Suskauer et al., 2008a; Suskauer et al., 2008b).

Thus, one interpretation of our results is that the FAT is involved in attention per se, and not necessarily inhibitory control or conflict detection. Indeed, one critique of the notion that the right IFG is associated with inhibition is that the typical experimental paradigms employed are assessing attentional processes (Hampshire et al., 2010; Chatham et al., 2012). For example, Chatham and colleagues (Chatham et al., 2012) and Hampshire and colleagues (Hampshire et al., 2010; Erika-Florence et al., 2014; Hampshire, 2015) have suggested that so-called “inhibitory control tasks” really tap into controlled context-monitoring processes, not inhibition. The authors further suggested that impairments in context-monitoring, supported by right IFG and associated circuits, might explain deficits seen in ADHD. They pointed to increased reaction time variability in SST paradigms in people with ADHD as support for such a contention (Castellanos et al., 2006), and suggest that treatments focusing on improving context-monitoring, rather than improving inhibitory control, might be more appropriately targeting the underlying deficit in ADHD. But these tasks confound context monitoring, conflict detection, and inhibitory control processes proposed to recruit the right IFG (Hampshire, 2015). Although some attempts have been made to tease these processes apart (Erika-Florence et al., 2014; Hampshire, 2015), there is still debate about whether right IFG is involved in attention more generally (Ridderinkhof et al., 2004), or more specifically inhibitory control (Aron et al., 2014).

It has also been proposed that a primary deficit in ADHD is in fact one of inhibitory control (Barkley, 1997; Schachar et al., 2000; Neely et al., 2017). However, inhibitory control and more broadly defined executive function deficits are not a universal feature of ADHD (Nigg et al., 2005), and in fact there may be executive function subtypes of ADHD, with an inhibitory control dysfunction profile describing only one of the subtypes (Roberts et al., 2017). These subtypes are defined at the behavioral level, and further progress in demarcating them may require the additional of data at other levels of analysis, such as at the neurobiological level. In this case, our findings suggest that delineation of the FAT in people with ADHD, and exploration of its functional relationship to executive function, might be important for understanding and dissociating ADHD subtypes. Indeed, our data reinforce the notion that attention problems associated with the FAT are explained by individual differences in executive function. However, our sample is a typical sample, and does not speak to whether there are subtypes that might be apparent in a clinical population. This would require further work in clinical populations, such as people diagnosed with ADHD.

### Limitations

One potential limitation of our study is the use of behavior report measures as a proxy for executive function and attention. This is a legitimate criticism, and the study should be in part viewed as a point of departure for future detailed investigations using laboratory paradigms. However, the behavioral ratings we used here have substantial construct validity and reliability (Achenbach and Rescorla, 2000; Gioia et al., 2000; Achenbach and Rescorla, 2001, 2003), and provide information that cannot necessarily be obtained from laboratory tasks. For example, Barkley and colleagues (Barkley and Murphy, 2010; Barkley and Fischer, 2011) found that ratings of executive function can sometimes be a better predictor of everyday impairment than laboratory tests of executive function. Rating scales are also effective with preschool children and perform as well as laboratory tasks, such as continuous performance tasks, at differentiating children with ADHD from typical children (Cak et al., 2017). Thus, while future research should incorporate laboratory tasks, it does not discount the utility of the results we report here.

### Conclusions

The work we report here shows that the FAT develops in a protracted manner into late adolescence/early adulthood, and that right lateralization of the fiber pathway is significantly associated with executive function. This fits with the putative functional roles of the regions the pathway connects—the right IFG and right pre-SMA. These results suggest that the FAT should be explored more carefully in research on developing executive function, or dysfunction as occurs in externalizing disorders such as ADHD.

## Acknowledgements

This work was supported by grants from the National Institutes of Mental Health (Grant R56MH108616 to P.G. and A.S.D) and from the National Institute for Drug Abuse (U01DA041156; salary support to A.S.D.).

## References

Achenbach TM, Rescorla L (2000) Manual for the ASEBA preschool forms & profiles: an integrated system of multi-informant assessment. Burlington, VT: ASEBA.

Achenbach TM, Rescorla L (2001) Manual for the ASEBA school-age forms & profiles: an integrated system of multi-informant assessment. Burlington, VT: ASEBA.

Achenbach TM, Rescorla L (2003) Manual for the ASEBA adult forms & profiles: for ages 18-59: adult self-report, adult behavior checklist. Burlington, VT: ASEBA.

Amador N, Fried I (2004) Single-neuron activity in the human supplementary motor area underlying preparation for action. J Neurosurg 100:250–259.

Aron AR, Robbins TW, Poldrack RA (2014) Inhibition and the right inferior frontal cortex: one decade on. Trends Cogn Sci 18:177–185.

Aron AR, Behrens TE, Smith S, Frank MJ, Poldrack RA (2007) Triangulating a cognitive control network using diffusion-weighted magnetic resonance imaging (MRI) and functional MRI. J Neurosci 27:3743–3752.

Aron AR, Herz DM, Brown P, Forstmann BU, Zaghloul K (2016) Frontosubthalamic Circuits for Control of Action and Cognition. J Neurosci 36:11489–11495.

Barkley RA (1997) Behavioral inhibition, sustained attention, and executive functions: constructing a unifying theory of ADHD. Psychol Bull 121:65–94.

Barkley RA, Murphy KR (2010) Impairment in occupational functioning and adult ADHD: the predictive utility of executive function (EF) ratings versus EF tests. Arch Clin Neuropsychol 25:157–173.

Barkley RA, Fischer M (2011) Predicting impairment in major life activities and occupational functioning in hyperactive children as adults: self-reported executive function (EF) deficits versus EF tests. Dev Neuropsychol 36:137–161.

Basilakos A, Fillmore PT, Rorden C, Guo D, Bonilha L, Fridriksson J (2014) Regional white matter damage predicts speech fluency in chronic post-stroke aphasia. Front Hum Neurosci 8:845.

Biederman J, Faraone SV, Doyle A, Lehman BK, Kraus I, Perrin J, Tsuang MT (1993) Convergence of the Child Behavior Checklist with structured interview-based psychiatric diagnoses of ADHD children with and without comorbidity. J Child Psychol Psychiatry 34:1241–1251.

Bozkurt B, Yagmurlu K, Middlebrooks EH, Karadag A, Ovalioglu TC, Jagadeesan B, Sandhu G, Tanriover N, Grande AW (2016) Microsurgical and Tractographic Anatomy of the Supplementary Motor Area Complex in Humans. World Neurosurg 95:99–107.

Broce I, Bernal B, Altman N, Tremblay P, Dick AS (2015) Fiber tracking of the frontal aslant tract and subcomponents of the arcuate fasciculus in 5-8-year-olds: Relation to speech and language function. Brain Lang 149:66–76.

Cak HT, Cengel Kultur SE, Gokler B, Oktem F, Taskiran C (2017) The Behavior Rating Inventory of Executive Function and Continuous Performance Test in Preschoolers with Attention Deficit Hyperactivity Disorder. Psychiatry Investig 14:260–270.

Carter AS, Briggs-Gowan MJ, Davis NO (2004) Assessment of young children’s social-emotional development and psychopathology: recent advances and recommendations for practice. Journal of Child Psychology and Psychiatry 45:109–134.

Castellanos FX, Sonuga-Barke EJ, Milham MP, Tannock R (2006) Characterizing cognition in ADHD: beyond executive dysfunction. Trends Cogn Sci 10:117–123.

Catani M, Dell’acqua F, Vergani F, Malik F, Hodge H, Roy P, Valabregue R, Thiebaut de Schotten M (2012) Short frontal lobe connections of the human brain. Cortex 48:273–291.

Catani M, Mesulam MM, Jakobsen E, Malik F, Martersteck A, Wieneke C, Thompson CK, Thiebaut de Schotten M, Dell’Acqua F, Weintraub S, Rogalski E (2013) A novel frontal pathway underlies verbal fluency in primary progressive aphasia. Brain 136:2619–2628.

Chatham CH, Claus ED, Kim A, Curran T, Banich MT, Munakata Y (2012) Cognitive control reflects context monitoring, not motoric stopping, in response inhibition. PLoS One 7:e31546.

Coxon JP, Van Impe A, Wenderoth N, Swinnen SP (2012) Aging and inhibitory control of action: cortico-subthalamic connection strength predicts stopping performance. J Neurosci 32:8401–8412.

Daducci A et al. (2014) Quantitative comparison of reconstruction methods for intra-voxel fiber recovery from diffusion MRI. IEEE Trans Med Imaging 33:384–399.

Descoteaux M, Angelino E, Fitzgibbons S, Deriche R (2007) Regularized, fast, and robust analytical Q-ball imaging. Magn Reson Med 58:497–510.

Desikan RS, Segonne F, Fischl B, Quinn BT, Dickerson BC, Blacker D, Buckner RL, Dale AM, Maguire RP, Hyman BT, Albert MS, Killiany RJ (2006) An automated labeling system for subdividing the human cerebral cortex on MRI scans into gyral based regions of interest. Neuroimage 31:968–980.

Duann JR, Ide JS, Luo X, Li CS (2009) Functional connectivity delineates distinct roles of the inferior frontal cortex and presupplementary motor area in stop signal inhibition. J Neurosci 29:10171–10179.

Duvernoy HM, Bourgouin P, Cabanis EA, Cattin F, Guyot J, Iba-Zizen MT, Maeder P, Parratte B, Tatu L, Vuillier F, vannson JL (1999) The human brain: Surface, three-dimensional sectional anatomy with MRI, and blood supply. New York: Springer-Wien.

Erika-Florence M, Leech R, Hampshire A (2014) A functional network perspective on response inhibition and attentional control. Nat Commun 5:4073.

Everts R, Lidzba K, Wilke M, Kiefer C, Mordasini M, Schroth G, Perrig W, Steinlin M (2009) Strengthening of laterality of verbal and visuospatial functions during childhood and adolescence. Hum Brain Mapp 30:473–483.

Favre E, Ballanger B, Thobois S, Broussolle E, Boulinguez P (2013) Deep brain stimulation of the subthalamic nucleus, but not dopaminergic medication, improves proactive inhibitory control of movement initiation in Parkinson’s disease. Neurotherapeutics 10:154–167.

Fujii M, Maesawa S, Motomura K, Futamura M, Hayashi Y, Koba I, Wakabayashi T (2015) Intraoperative subcortical mapping of a language-associated deep frontal tract connecting the superior frontal gyrus to Broca’s area in the dominant hemisphere of patients with glioma. J Neurosurg 122:1390–1396.

Furniss T, Beyer T, Guggenmos J (2006) Prevalence of behavioural and emotional problems among six-years-old preschool children. Social Psychiatry and Psychiatric Epidemiology 41:394–399.

Garavan H, Ross TJ, Kaufman J, Stein EA (2003) A midline dissociation between error-processing and response-conflict monitoring. Neuroimage 20:1132–1139.

Gioia GA, Isquith PK, Guy SC, Kenworthy L (2000) Behavior Rating Inventory of Executive Function. Lutz, FL: Psychological Assessment Resources.

Gioia GA, Isquith PK, Retzlaff PD, Espy KA (2002) Confirmatory factor analysis of the Behavior Rating Inventory of Executive Function (BRIEF) in a clinical sample. Child Neuropsychol 8:249–257.

Hampshire A (2015) Putting the brakes on inhibitory models of frontal lobe function. Neuroimage 113:340–355.

Hampshire A, Chamberlain SR, Monti MM, Duncan J, Owen AM (2010) The role of the right inferior frontal gyrus: inhibition and attentional control. Neuroimage 50:1313–1319.

Hayes AF (2013) Introduction to mediation, moderation, and conditional process analysis: a regression-based approach. New York: The Guilford Press.

Herve PY, Zago L, Petit L, Mazoyer B, Tzourio-Mazoyer N (2013) Revisiting human hemispheric specialization with neuroimaging. Trends Cogn Sci 17:69–80.

Holland SK, Plante E, Weber Byars A, Strawsburg RH, Schmithorst VJ, Ball WS, Jr. (2001) Normal fMRI brain activation patterns in children performing a verb generation task. Neuroimage 14:837–843.

Isquith PK, Gioia GA (2000) BRIEF predictions of ADHD: clinical utility of the Behavior Rating Inventory of Executive Function for detecting ADHD subtypes in children. Arch Clin Neuropsych 15:780–781.

Jahanshahi M (2013) Effects of deep brain stimulation of the subthalamic nucleus on inhibitory and executive control over prepotent responses in Parkinson’s disease. Front Syst Neurosci 7:118.

Jahanshahi M, Obeso I, Rothwell JC, Obeso JA (2015) A fronto-striato-subthalamic-pallidal network for goal-directed and habitual inhibition. Nat Rev Neurosci 16:719–732.

Jensen PS, Arnold LE, Swanson JM, Vitiello B, Abikoff HB, Greenhill LL, Hechtman L, Hinshaw SP, Pelham WE, Wells KC (2007) 3-year follow-up of the NIMH MTA study. Journal of the American Academy of Child & Adolescent Psychiatry 46:989–1002.

Keenan K, Wakschlag LS (2000) More than the terrible twos: The nature and severity of behavior problems in clinic-referred preschool children. J Abnorm Child Psych 28:33–46.

Kemerdere R, de Champfleur NM, Deverdun J, Cochereau J, Moritz-Gasser S, Herbet G, Duffau H (2016) Role of the left frontal aslant tract in stuttering: a brain stimulation and tractographic study. J Neurol 263:157–167.

Kim JH, Lee JM, Jo HJ, Kim SH, Lee JH, Kim ST, Seo SW, Cox RW, Na DL, Kim SI, Saad ZS (2010) Defining functional SMA and pre-SMA subregions in human MFC using resting state fMRI: functional connectivity-based parcellation method. Neuroimage 49:2375–2386.

Kinoshita M, de Champfleur NM, Deverdun J, Moritz-Gasser S, Herbet G, Duffau H (2015) Role of fronto-striatal tract and frontal aslant tract in movement and speech: an axonal mapping study. Brain Struct Funct 220:3399–3412.

Kinoshita M, Shinohara H, Hori O, Ozaki N, Ueda F, Nakada M, Hamada J, Hayashi Y (2012) Association fibers connecting the Broca center and the lateral superior frontal gyrus: a microsurgical and tractographic anatomy. J Neurosurg 116:323–330.

Kronfeld-Duenias V, Amir O, Ezrati-Vinacour R, Civier O, Ben-Shachar M (2016) The frontal aslant tract underlies speech fluency in persistent developmental stuttering. Brain Struct Funct 221:365–381.

Madsen KS, Baare WF, Vestergaard M, Skimminge A, Ejersbo LR, Ramsoy TZ, Gerlach C, Akeson P, Paulson OB, Jernigan TL (2010) Response inhibition is associated with white matter microstructure in children. Neuropsychologia 48:854–862.

Mandelli ML, Caverzasi E, Binney RJ, Henry ML, Lobach I, Block N, Amirbekian B, Dronkers N, Miller BL, Henry RG, Gorno-Tempini ML (2014) Frontal white matter tracts sustaining speech production in primary progressive aphasia. J Neurosci 34:9754–9767.

Martino J, De Lucas EM (2014) Subcortical anatomy of the lateral association fascicles of the brain: A review. Clin Anat 27:563–569.

McCandless S, L OL (2007) The Clinical Utility of the Behavior Rating Inventory of Executive Function (BRIEF) in the diagnosis of ADHD. J Atten Disord 10:381–389.

Molina BS, Hinshaw SP, Swanson JM, Arnold LE, Vitiello B, Jensen PS, Epstein JN, Hoza B, Hechtman L, Abikoff HB (2009) The MTA at 8 years: prospective follow-up of children treated for combined-type ADHD in a multisite study. Journal of the American Academy of Child & Adolescent Psychiatry 48:484–500.

Molina BS, Hinshaw SP, Eugene Arnold L, Swanson JM, Pelham WE, Hechtman L, Hoza B, Epstein JN, Wigal T, Abikoff HB (2013) Adolescent substance use in the multimodal treatment study of attention-deficit/hyperactivity disorder (ADHD)(MTA) as a function of childhood ADHD, random assignment to childhood treatments, and subsequent medication. Journal of the American Academy of Child & Adolescent Psychiatry 52:250–263.

Mostofsky SH, Cooper KL, Kates WR, Denckla MB, Kaufmann WE (2002) Smaller prefrontal and premotor volumes in boys with attention-deficit/hyperactivity disorder. Biol Psychiatry 52:785–794.

Nachev P, Wydell H, O’Neill K, Husain M, Kennard C (2007) The role of the pre-supplementary motor area in the control of action. Neuroimage 36 Suppl 2:T155–163.

Neely KA, Wang P, Chennavasin AP, Samimy S, Tucker J, Merida A, Perez-Edgar K, Huang-Pollock C (2017) Deficits in inhibitory force control in young adults with ADHD. Neuropsychologia 99:172–178.

Nigg JT, Willcutt EG, Doyle AE, Sonuga-Barke EJ (2005) Causal heterogeneity in attention-deficit/hyperactivity disorder: do we need neuropsychologically impaired subtypes? Biol Psychiatry 57:1224–1230.

Obeso I, Wilkinson L, Casabona E, Speekenbrink M, Luisa Bringas M, Alvarez M, Alvarez L, Pavon N, Rodriguez-Oroz MC, Macias R, Obeso JA, Jahanshahi M (2014) The subthalamic nucleus and inhibitory control: impact of subthalamotomy in Parkinson’s disease. Brain 137:1470–1480.

Papachristou E, Schulz K, Newcorn J, Bedard AC, Halperin JM, Frangou S (2016) Comparative Evaluation of Child Behavior Checklist-Derived Scales in Children Clinically Referred for Emotional and Behavioral Dysregulation. Front Psychiatry 7:146.

Ridderinkhof KR, Ullsperger M, Crone EA, Nieuwenhuis S (2004) The role of the medial frontal cortex in cognitive control. Science 306:443–447.

Roberts BA, Martel MM, Nigg JT (2017) Are There Executive Dysfunction Subtypes Within ADHD? J Atten Disord 21:284–293.

Rubia K, Overmeyer S, Taylor E, Brammer M, Williams SC, Simmons A, Bullmore ET (1999) Hypofrontality in attention deficit hyperactivity disorder during higher-order motor control: a study with functional MRI. Am J Psychiatry 156:891–896.

Schachar R, Mota VL, Logan GD, Tannock R, Klim P (2000) Confirmation of an inhibitory control deficit in attention-deficit/hyperactivity disorder. J Abnorm Child Psychol 28:227–235.

Sierpowska J, Gabarros A, Fernandez-Coello A, Camins A, Castaner S, Juncadella M, de Diego-Balaguer R, Rodriguez-Fornells A (2015) Morphological derivation overflow as a result of disruption of the left frontal aslant white matter tract. Brain Lang 142:54–64.

Sonuga-Barke EJ, Brandeis D, Cortese S, Daley D, Ferrin M, Holtmann M, Stevenson J, Danckaerts M, Van der Oord S, Döpfner M (2013) Nonpharmacological interventions for ADHD: systematic review and meta-analyses of randomized controlled trials of dietary and psychological treatments. Am J Psychiat 170:275–289.

Suskauer SJ, Simmonds DJ, Caffo BS, Denckla MB, Pekar JJ, Mostofsky SH (2008a) fMRI of intrasubject variability in ADHD: anomalous premotor activity with prefrontal compensation. J Am Acad Child Adolesc Psychiatry 47:1141–1150.

Suskauer SJ, Simmonds DJ, Fotedar S, Blankner JG, Pekar JJ, Denckla MB, Mostofsky SH (2008b) Functional magnetic resonance imaging evidence for abnormalities in response selection in attention deficit hyperactivity disorder: differences in activation associated with response inhibition but not habitual motor response. J Cogn Neurosci 20:478–493.

Swann NC, Cai W, Conner CR, Pieters TA, Claffey MP, George JS, Aron AR, Tandon N (2012) Roles for the pre-supplementary motor area and the right inferior frontal gyrus in stopping action: electrophysiological responses and functional and structural connectivity. Neuroimage 59:2860–2870.

Szmuda T, Rogowska M, Sloniewski P, Abuhaimed A, Szmuda M, Springer J, Sabisz A, Dzierzanowski J, Starzynska A, Przewozny T, Skorek A (2017) Frontal aslant tract projections to the inferior frontal gyrus. Folia Morphol (Warsz).

Thomas JM, Guskin KA (2001) Disruptive behavior in young children: what does it mean? J Am Acad Child Adolesc Psychiatry 40:44–51.

Toga AW, Thompson PM (2003) Mapping brain asymmetry. Nat Rev Neurosci 4:37–48.

Tournier JD, Calamante F, Connelly A (2007) Robust determination of the fibre orientation distribution in diffusion MRI: non-negativity constrained super-resolved spherical deconvolution. Neuroimage 35:1459–1472.

Tournier JD, Calamante F, Connelly A (2013) Determination of the appropriate b value and number of gradient directions for high-angular-resolution diffusion-weighted imaging. NMR Biomed 26:1775–1786.

van Wouwe NC, Pallavaram S, Phibbs FT, Martinez-Ramirez D, Neimat JS, Dawant BM, D’Haese PF, Kanoff KE, van den Wildenberg WPM, Okun MS, Wylie SA (2017) Focused stimulation of dorsal subthalamic nucleus improves reactive inhibitory control of action impulses. Neuropsychologia 99:37–47.

Vassal F, Boutet C, Lemaire JJ, Nuti C (2014) New insights into the functional significance of the frontal aslant tract: an anatomo-functional study using intraoperative electrical stimulations combined with diffusion tensor imaging-based fiber tracking. Br J Neurosurg 28:685–687.

Vergani F, Lacerda L, Martino J, Attems J, Morris C, Mitchell P, Thiebaut de Schotten M, Dell’Acqua F (2014) White matter connections of the supplementary motor area in humans. J Neurol Neurosurg Psychiatry 85:1377–1385.

Wiecki TV, Frank MJ (2013) A computational model of inhibitory control in frontal cortex and basal ganglia. Psychol Rev 120:329–355.

Yeh FC, Wedeen VJ, Tseng WY (2010) Generalized q-sampling imaging. IEEE Trans Med Imaging 29:1626–1635.

Yeh FC, Verstynen TD, Wang Y, Fernandez-Miranda JC, Tseng WY (2013) Deterministic diffusion fiber tracking improved by quantitative anisotropy. PLoS One 8:e80713.

